# *In vivo* base editing reduces liver cysts in autosomal dominant polycystic kidney disease

**DOI:** 10.1101/2025.02.07.636600

**Authors:** Antonia Ibel, Rishi Bhardwaj, Duygu Elif Yilmaz, Shuhan Kong, Sarah Wendlinger, Dimitra Papaioannou, Claudia Diezemann, Kai-Uwe Eckardt, Fatima Hasan, Verena Klämbt, Jan Halbritter, Sorin Fedeles, Matteus Krappitz, Michael M. Kaminski

**Affiliations:** Department of Nephrology and Medical Intensive Care, Charité Universitätsmedizin, Berlin, Germany; BIMSB, Max Delbrück Center for Molecular Medicine in the Helmholtz Association, Berlin, Germany; Department of Internal Medicine, Section of Nephrology, Yale School of Medicine, New Haven, Connecticut, USA; Department of Pediatric Gastroenterology, Nephrology and Metabolic Diseases, Charité Universitätsmedizin, Berlin, Germany; Berlin Institute of Health, BIH Charité Clinician Scientist Program, Berlin, Germany; Division of Renal Diseases and Hypertension, School of Medicine, CU Anschutz, University of Colorado, Aurora, Colorado, USA

## Abstract

Autosomal dominant polycystic kidney disease (ADPKD) is the most prevalent genetic kidney disorder, affecting over 10 million individuals worldwide. Cystic expansion typically progresses to kidney failure and also involves the liver with limited treatment options. Pathogenic variants in *PKD1* or *PKD2* account for 85-90% of cases. Genetic re-expression of *Pkd1* or *Pkd2* has been shown to partially reverse key characteristics of the disease phenotype in mice. Despite advancements in the understanding of the genetic basis, it remains unclear whether the correction of underlying pathogenic variants can effectively prevent, modify, or reverse the disease. Additionally, the feasibility of extrinsically delivered genome editing as a treatment option for ADPKD remains largely unexplored. In this study, we employed CRISPR base editing to correct a spectrum of representative pathogenic *PKD1* variants selected from a patient cohort achieving precise and efficient editing *in vitro*. Correction of a representative murine missense variant (c.6646C>T (R2216W)) in primary renal epithelial cells successfully increased polycystin-1 expression and reduced levels of the endoplasmic reticulum stress marker sXBP1. *In vivo*, base editor delivery to the c.6646C>T (R2216W) knock-in mouse enabled correction of the pathogenic variant, resulting in a significant reduction in liver cysts. These findings provide the first evidence of ADPKD reversibility through genome editing, opening promising novel therapeutic perspectives for affected patients and their families.

## Introduction

Autosomal dominant polycystic kidney disease (ADPKD) is the most prevalent genetic cause of chronic kidney disease, with a prevalence between 1 in 400 and 1 in 1,000 individuals^1,2^. ADPKD affects over 10 million people globally and thus represents a significant health burden. It is a multisystemic disorder characterized by polycystic kidney and liver disease (PLD) along with additional extrarenal manifestations such as intracranial arterial aneurysms. Renal cysts originate from multiple tubular segments of the nephron, gradually expanding over lifetime. This growth compresses surrounding renal tubules, leading to cystic kidney enlargement, inflammation, fibrosis, and progression to kidney failure, typically occurring between the ages of 40 and 70^1,2^. Common complications of liver or kidney cysts include compression of the neighboring intraperitoneal and intrathoracic organs, infections and pain. Tolvaptan, a vasopressin V2 receptor antagonist, is the only approved therapy to slow disease progression, albeit with limited efficacy and severe side effects. Importantly, Tolvaptan does not affect liver cysts, leaving patients with progressive liver cysts without any pharmacological treatment option. As at least 10% of ADPKD patients suffer from symptomatic PLD, there is an urgent medical need for developing treatment options.

Around 85-90% of ADPKD cases are due to variants in *PKD1* (75%) or *PKD2* (10-15%). Patients with *PKD1* variants generally exhibit an earlier onset and more severe disease progression. *PKD1* and *PKD2* encode polycystin-1 (PC1) and polycystin-2 (PC2), respectively, key components of a calcium-permeable ion channel in renal tubular cells that are crucial for intracellular signaling in primary cilia. Additionally, pathogenic variants in genes such as *IFT140, GANAB, ALG5, ALG8, ALG9*, or *DNAJB11* account for less than 1% of ADPKD-like phenotypes. ADPKD patients typically carry a heterozygous *PKD1* or *PKD2* germline mutation, and cyst formation and disease progression often require a “second-hit”, which may involve somatic inactivation of the wild type *PKD1* or *PKD2* allele, variants in other ADPKD-related genes, environmental factors, or unidentified genetic modifiers. Additionally, it has been suggested that cystogenesis in ADPKD is influenced by gene dosage thresholds, with disease severity correlating to functional levels of polycystin proteins^3,4^. Consequently, loss-of-function variants are associated with more severe phenotypes, whereas milder, late-onset forms of the disease are linked to hypomorphic missense mutations that partially preserve polycystin function^1,5^. Studies have demonstrated that reduced levels of PC1 in animal models are sufficient to induce renal cyst formation. Conversely, genetic reactivation of *Pkd1* or *Pkd2* in murine ADPKD mouse models results in a partial reversal of the phenotype^6^. Additionally, the genetic deletion or pharmaceutical inhibition of the miR-17 motif within the 3’ UTR of the *PKD1* or *PKD2* genes, which increases PC1 and PC2 levels, attenuated renal cyst growth in an experimental *Pkd1*-mutant mouse model, even after disease onset^7^. While re-expression of *Pkd1* or *Pkd2* in mouse models can reverse certain aspects of ADPKD, even in advanced stages^6–8^, it is unknown whether direct correction of pathogenic *PKD1* variants through genome editing may similarly restore *PKD1* function. Base editors are CRISPR/Cas-based genome editing tools that enable the direct conversion of single bases without the need of double strand DNA breaks, DNA templates or homology directed repair^9,10^. They consist of a catalytic disabled Cas enzyme fused to a deaminase, enabling a C-to-T^9^, A-to-G^10^, A-to-Y^11^, and C-to-G^12^ edit. Recent preclinical studies have highlighted their potential and safety profile for *in vivo* genome editing in various genetic diseases^13–16^. However, no genome editing approach has been applied to prevent, halt or reverse ADPKD.

Here, we explore base editing as a potential treatment of ADPKD by focusing on its most important extrarenal complication: polycystic liver disease.

## Methods

### Cloning strategy and plasmids

SgRNA sequences were synthesized as dsDNA fragments (Eurofins Genomics). The duplexed oligos of the corresponding spacer sequence were annealed and ligated into the BsmBI-digested pU6-pegRNA-GG-acceptor plasmid provided by David Liu (Addgene plasmid #132777). pCMV_BE4max, pCMV_ABEmax, and ABE8e were a gift from David Liu (Addgene plasmids #112093, #112095 and #138489). ABE8.20-m was a gift from Nicole Gaudelli (Addgene plasmid # 136300). pCMV-T7-ABE8e-nSpG-P2A-EGFP (KAC984), pCAG-CBE4max-SpG-P2A-EGFP (RTW4552), and pCMV-T7-ABE8e-nSpRY-P2A-EGFP (KAC1069) were a gift from Benjamin Kleinstiver (Addgene plasmids #185911, #139998, and #185912)^17–22^.

### Cell culture, transfection and DNA isolation

HEK293T cell lines were maintained in DMEM (Thermo Fisher Scientific) supplemented with 10% (v/v) fetal bovine serum (Sigma F7524) and 1% P/S at 37 °C with 5% CO_2_. HEK293T cell lines were seeded at low passages with a density of 5^*^10^5^ cells per 300 µl in antibiotic-free medium in a 48-well cell culture microplate (Falcon CLS351172) and grown for 24 hours at 37 °C with 5% CO_2_. Transfection with sgRNA and base editor plasmids was performed using the TransIT-X2 Dynamic Delivery System (Mirus Bio MIR 6004) following manufacturer’s protocol in a 3:1 BE:sgRNA ratio by weight. 260 ng total plasmid was transfected per well. 72 hours after transfection, genomic DNA (gDNA) isolation was performed. For HEK293T cell line gDNA isolation, cells were washed once with PBS (DPBS, Gibco 14190250), trypsinized, spun down at 250 g for 5 minutes. Pellets were resuspended in 100 µl lysis buffer and incubated for 16 hours at 55 °C on a heat block at 300 rpm. gDNA was isolated and purified using MagBinding Beads (Zymo Research D4100-2-24) at a 1x ratio, with three washing steps using 80% Ethanol and eluted in EB buffer. MagBinding Beads were prepared according to manufacturer’s protocol. For quantification of gDNA, the Quant-iT dsDNA HS assay (Invitrogen Q33232) was used.

Renal tubular epithelial cells (RTECs) were maintained in DMEM/F12 (Thermo Fisher Scientific) supplemented with 10% (v/v) fetal bovine serum (Sigma F7524) and 1% P/S at 37 °C with 5% CO_2_. RTECs were seeded at low passages with a density of 4×10^5^ cells per 2 mL in antibiotic-free medium in a 6-well cell culture microplate (Falcon CLS351172) and grown for 24 hours at 37 °C with 5% CO_2_. Transfection with sgRNA and base editor plasmids was performed using the TransIT-X2 Dynamic Delivery System (Mirus Bio MIR 6004) following manufacturer’s protocol in a 3:1 BE:sgRNA ratio by weight. 2.5 µg total plasmid was transfected per well. 72 hours after transfection they were single-cell sorted for GFP expression and colonies of GFP positive cells were expanded and sequenced. RTEC gDNA isolation was performed as mentioned above.

### Targeted Amplicon Sequencing

Next-generation targeted amplicon sequencing (TAS) was performed to determine the efficiency of genome modification at the target sites using a two-step PCR-based library construction method adapted from the Illumina Nextera XT DNA library preparation. The target loci were amplified from 100 ng of gDNA using the Q5 Hot Start high Fidelity 2x MM (NEB) and PCR-1 primers (Suppl. Table 1). Primers for the engineered HEK293T cell lines were used according to the respective construct or designed with PrimerBlast by NCBI with overhangs allowing for the subsequent indexing PCR. The PCR products were purified using MagBinding Beads (Zymo Research D4100-2-24) at a 0.8x ratio and quantified using the Quant-iT dsDNA HS assay. Around 20 ng of purified PCR-1 products was used as template for the indexing PCR (PCR-2) to add barcodes and Illumina adapter sequences using Q5 and primers (Suppl. Table 1). PCR products were again purified using MagBinding Beads at 0.7x ratio, quantified and pooled equimolar. Pooled libraries were checked with the D1000 ScreenTape system (Agilent), spiked with 30-60% PhiX (Illumina), depending on the library complexity, and subsequently denatured. The final library was loaded on a MiniSeq sequencer at 1.5 pM and sequenced using a Mini Seq Mid Output Kit (300 cycles) (Illumina). On-target genome editing efficiencies and bystander edits were determined from sequencing data using CRISPResso2^23^.

### Generation of HEK293T reporter cell line

HEK293T cell lines were engineered to carry pathogenic *PKD1* variants by using the sleeping beauty transposase system^24^. The variants, flanked by 150 bps of surrounding genomic sequence, including specific primer sequences on the 5’ and 3’ ends (Suppl. Table 1) for targeted sequencing, were synthesized as dsDNA fragments (IDT gBlocks). Using Gibson assembly, the DNA fragments were cloned into the pT4 SB plasmid (gift from Zsuzsanna Iszvak). HEK293T cells were electroporated with 4 µg pT4 transposon plasmid together with 1 µg of the SB100x transposase RNA using the 4D-Nucleofector (Lonza). After culturing the cells for 3 days they were single-cell sorted for low GFP expression and colonies of single clones expanded and sequenced for the correct integration of targets.

### AAV production

Cbh_v5_AAV-ABE_N and Cbh_v5_AAV-ABE_C were a gift from David Liu (Addgene plasmids #137177 & #137178) with the C-terminal part carrying sgRNA_45^25^. The duplexed oligos of the corresponding spacer sequence were annealed and ligated into the BsmBI-digested C-terminal plasmid. The AAV vectors used were constructed and packaged by the Charité Core Facility or VectorBuilder. Clean up was done using an iodixanol gradient centrifugation or cesium chloride; pAdDeltaF6 and pAAV2/8 were used as helper and REP/CAP plasmids.

### Murine experiments

The mice described in this study are on a C57BL6/129 mixed background. Mice of both sexes were used. The mouse lines used in this study were previously described and include *Pkd1*^*R2216W/fl* 26^ and *UBC-Cre-ERT2*^27^. The *Pkd1*^*R2216W/fl*^;*UBC-Cre-ERT2* mice display strong Cre expression in the liver bile ducts and kidney proximal tubules upon tamoxifen induction. We induced the deletion of the *Pkd1*-floxed allele with tamoxifen between P28 to P42 followed by dual AAV8 split-ABE intraperitoneal (i.p.) injection (one-time administration) at P49 with a total concentration of 8×10^11^ VG/mouse. The phenotype was analyzed 12 weeks post-treatment. We examined liver to body weight ratio, and liver cystic index. We used AAV8-GFP i.p. administration to confirm tissue/cell-type tropism. Animal numbers for each study were determined by power calculations before initiation of the study. All animals used in this study were in accordance with scientific, human and ethical principles and in compliance with animal welfare regulations approved by the Yale Institutional Animal Care and Use Committee.

### Protein Preparation and Immunoblot Analysis

Cultured cells were extracted and homogenized in an ice-cold homogenization buffer (250 mM sucrose, 10 mM triethanolamine, pH 8.45 containing protease inhibitors). The homogenates were then sonicated 5 times for 1 second each, followed by centrifugation at 1000 *g* for 10 minutes. The resulting supernatant was analyzed as total lysate. Immunoblotting was performed using rabbit anti-HSP90 (Santa Cruz Biotechnology; sc-7947; 1:5000), mouse anti-PC1, 7E12 (Invitrogen; mA5–15253; 1:500), rabbit anti-XBP1s (Abcam; ab220783; 1:2000), mouse anti-ß Actin (Sigma; A2228; 1:5000). Secondary antibodies included anti-mouse/rabbit HRP-conjugates (1:2000; Jackson ImmunoResearch Laboratories, West Grove, PA) and were incubated with the membrane for 1 hour at room temperature. Thermo Scientific Pierce ECL Plus Western Blotting-Substrate (Thermo Scientific; #11527271) or SuperSignal West Femto Chemiluminescent Substrate (Thermo Fisher Scientific; #34094) was used for chemiluminescence detection. The volume of individual immunoblot bands, in pixels, was determined by optical densitometry using ImageJ software (NIH).

### Liver Histological Assessment

Mice were anesthetized by injecting ketamine/xylazine intraperitoneally followed by cardiac perfusion with 1x PBS. The liver was then extracted, one part of which was snap frozen and the remaining liver was fixed in 10% formalin for histological sectioning (5 µm) at Research Histology Lab, Department of Comparative Medicine, Yale University. Hematoxylin-Eosin sections thus obtained were imaged and scanned (4x) to measure the cystic index using a Nikon Eclipse TE2000-U microscope using MetaMorph software (Universal Imaging)^28^.

### Statistical Analysis

Comparisons of three or more groups were performed using one-way ANOVA followed by Tukey’s multiple group comparison post-hoc test. Comparison of two groups was performed using the two-tailed *t* test. A *P* value of <0.05 was considered the threshold for statistical significance. Data are presented as the mean ± SD.

## Results

### Correction of pathogenic *PKD1* variants selected from an ADPKD patient cohort using adenine and cytosine base editing

Point mutations represent the largest category of human pathogenic variants, accounting for 58% of all pathogenic genetic changes^29^. Common base editors, such as adenine base editors (ABEs) and cytosine base editors (CBEs), could theoretically correct up to 61% of these mutations. To explore the applicability of base editing for pathogenic *PKD1* variants, we screened our local ADPKD cohort (**Fig. 1a**) and identified 39 representative variants distributed across exons 4 to 43. These included both missense and nonsense variants, which were potentially targetable with CBEs or ABEs (**Fig. 1b**). The variants were introduced into HEK293T cells using a transposase system, which stably integrated the pathogenic variant along with 150 bps of flanking *PKD1* genomic sequence (**Fig. 1a**). We tested different combinations of base editors and single guide RNAs (sgRNAs) with corresponding protospacer-adjacent motifs (PAMs) for optimal editing (**Fig. 1c**). While editing of only five variants showed correction efficiencies below 10%, editing of most variants achieved on-target efficiencies between 30% and 60%, comparable to the editing efficiency observed at a commonly used, highly efficient control site in the beta-2-microglobulin (*B2M*) gene (**Fig. 1c**). Notably, as part of our screening approach, we did not enrich for successfully transfected cells, likely leading to an underestimation of editing efficiency. These findings indicate that base editors can be broadly applied to correct pathogenic *PKD1* variants in ADPKD, including those affecting the most functionally relevant domains (e.g., REJ, PLAT), underscoring their clinical potential.

**Fig. 1:**
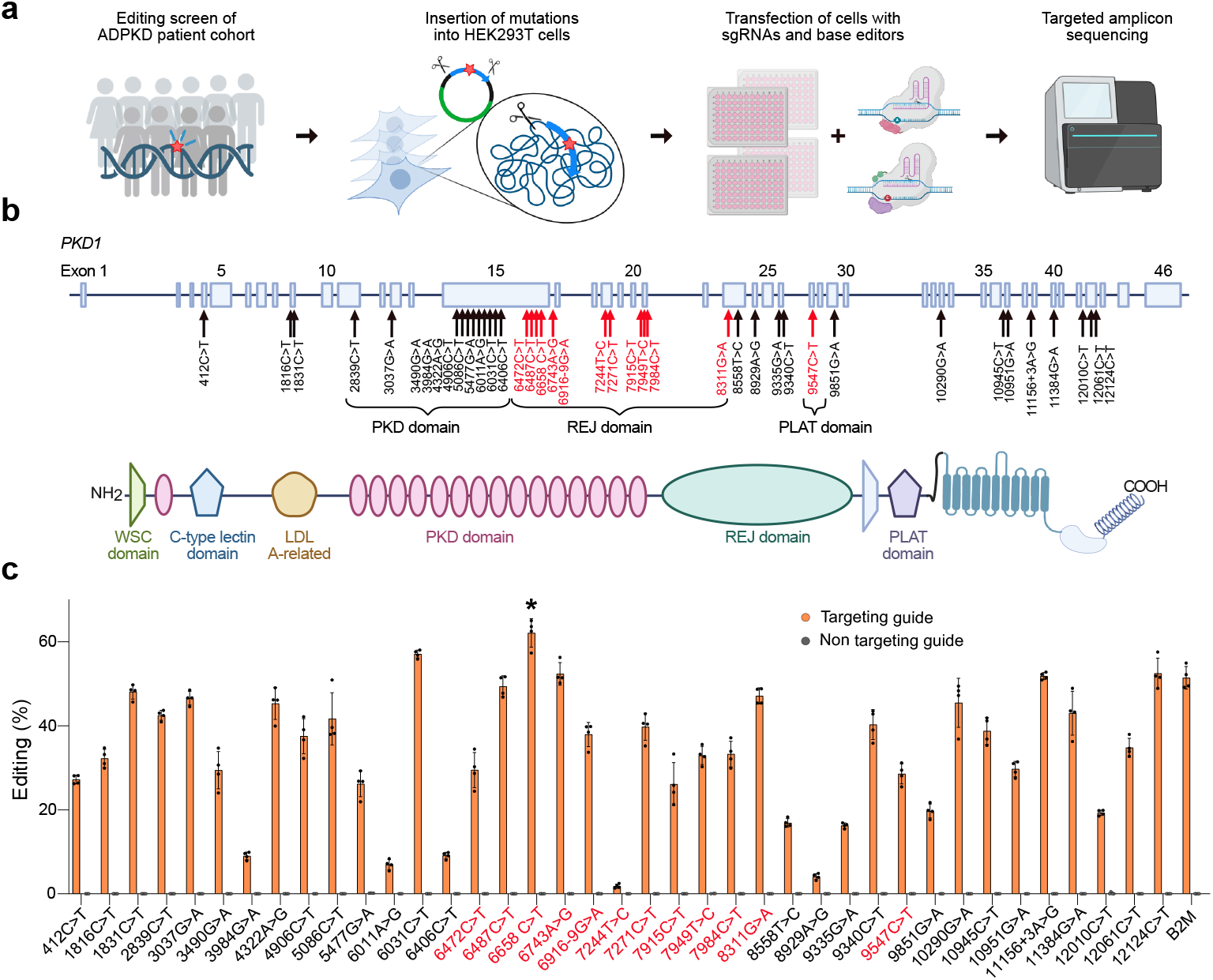
Base editing screen for correction of *PKD1* variants. **a,** Schematic indicating the experimental workflow; From 185 patients with typical ADPKD, 104 were selected with (likely) pathogenic single nucleotide variants, from which 39 variants correctable by adenine or cytosine base editors were introduced in HEK293T cells along with 150 bp of flanking endogenous sequence using a transposase system. Cells were then transfected with base editors and sgRNAs, followed by targeted amplicon sequencing. **b**, Genomic structure and protein domain structure of *PKD1*. Arrows indicate genomic positions of identified pathogenic variants from our ADPKD cohort, red indicates location in functionally important domains (e.g., REJ, PLAT). **c**, Editing efficiency across all different *PKD1* variants identified in our ADPKD cohort, tested in the engineered HEK293T cell line. Beta-2-microglobulin (*B2M*) served as a positive control. Asterisk highlights human c.6658C>T variant (R2220W, corresponding to murine R2216W) showing the highest editing efficiency. N = 4 biological replicates. Bar graphs indicate means ± SD.

To optimize on-target editing and reduce bystander editing, we selected the hypomorphic c.6658C>T (R2220W)^30^ pathogenic missense variant from our screen, for which a fully characterized knock-in mouse model is available mimicking key characteristics of human ADPKD. We tested three different base editors (ABEmax, ABE8.20m, ABE8e) with varying editing windows, in combination with suitable sgRNAs, to assess efficiency and precision for correction of the human c.6658C>T (R2220W) (**Fig. 2a**). The observed editing efficiencies for the intended A5>G conversion ranged from 10% to 65%, with ABE8e demonstrating the highest efficiency. We then assessed editing outcomes for potential unwanted bystander mutations and identified an edit at A10 (A10>G) and A13 (A13>G) within the editing window, resulting in a leucine to proline substitution at amino acid position p.2218 or valine to alanine substitution at amino acid position p.2217, respectively. This bystander edit was predominantly observed with ABE8e or ABE8.20m, while ABEmax achieved 20.9% ± 7.9% on-target editing efficiency without any bystander mutations and consequently was selected for further experiments. These findings demonstrate that the correction of c.6658C>T (R2220W) can be systematically optimized, ultimately yielding efficient and precise base editing using ABEmax.

**Fig. 2.**
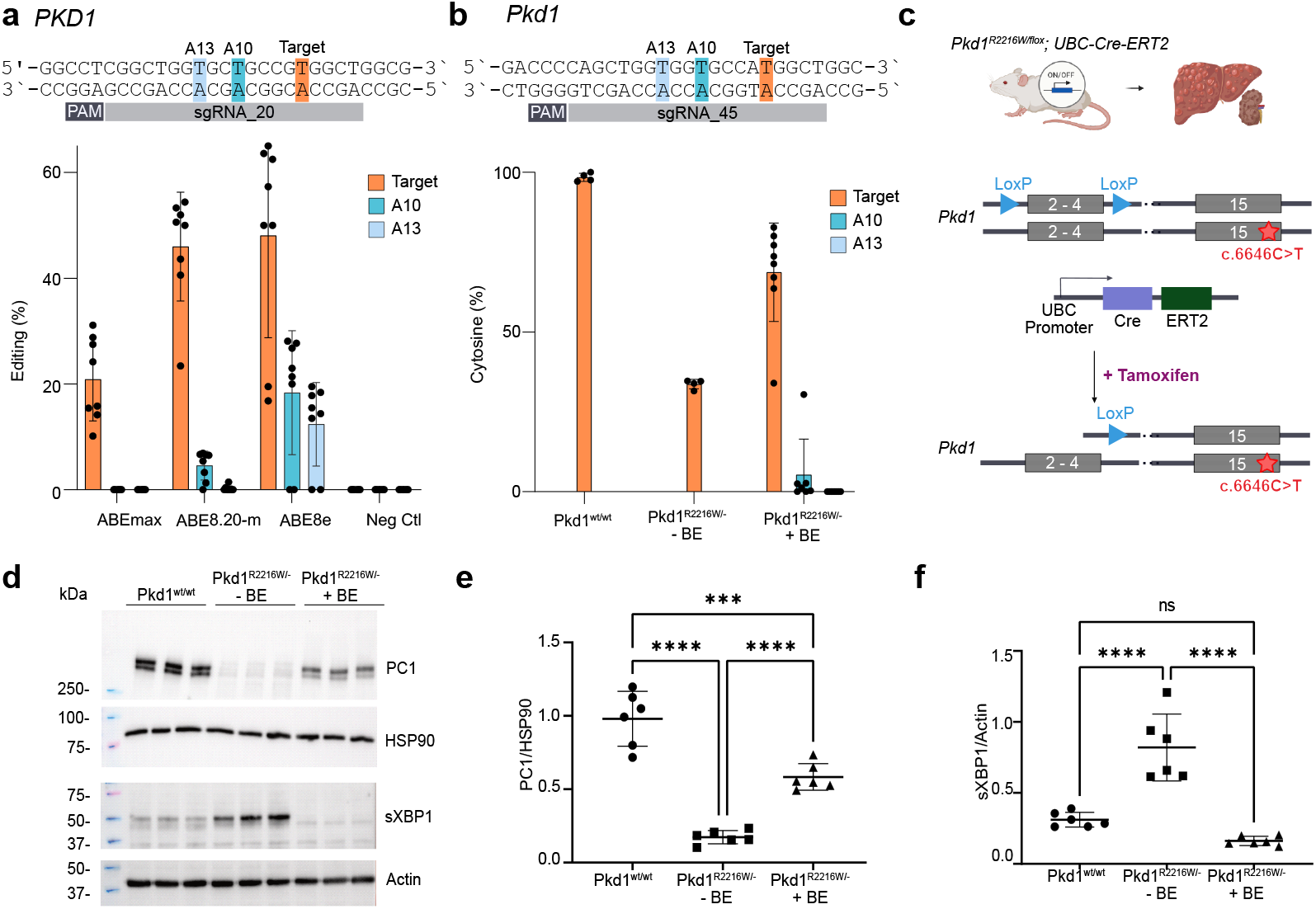
Correction and functional characterization of the RW missense variant *in vitro*. **a,** Schematic illustrating the targeting strategy; editing efficiency is shown for indicated base editors and sites in human *PKD1* c.6658C>T (R2220W). Orange indicates the intended on-target editing and blue indicates unintended bystander editing. **b**, Schematic illustrating the targeting strategy; editing efficiency is shown for indicated base editors and sites in corresponding mouse *Pkd1* c.6646C>T (R2216W). **c**, Schematic of the knock-in mouse model with the pathogenic *Pkd1* c.6646C>T (R2216W) on one allele and deactivation of the other allele through Cre-mediated excision of exons 2-4. UBC driven Cre expression upon tamoxifen administration leads to cystic liver disease and mild cystic kidney disease. **d**, Western blots indicating levels of the indicated proteins in base-edited cells and controls. sXBP1 served as a marker of ER stress. Representative immunoblots show signals for PC1 (NTF-fragment 450 kDa), spliced XBP1 (sXBP1; 56 kDa). **e**,**f**, Graphs represent the densitometric evaluation of PC1 and sXBP1; SHSP90 or Actin serves as loading controls (90 and 42 kDa, respectively) and are shown below the respective immunoblots. N = 3 biological replicates with two technical replicates. Data are the means ± SD, ∗∗∗p < 0.001, ∗∗∗∗p < 0.0001, ns: not significant.

### Functional characterization of *in vitro* base editing outcomes for correction of the murine *Pkd1* c.6646C>T (R2216W) variant

Next, we aimed to correct the corresponding mouse variant, c.6646C>T (R2216W) in primary renal epithelial cells (RTECs) derived from the *Pkd1*^*R2216W/-*^ knock-in mouse model (**Fig. 2b**). In these mice, the pathogenic variant is carried on one allele, while the other allele is deactivated through Cre-mediated excision of exons 2-4 upstream of the variant’s position, resulting in heterozygosity for the variant (**Fig. 2c**). ABEmax and sgRNA_45 yielded a mean cytosine recovery of 68.7% ± 15.4%, compared to 33.7% ± 1.5% in non-edited control cells, as determined by TAS (**Fig. 2b**). We observed no bystander editing at A13 (A13>G) and minimal bystander editing at A10 (A10>G) of 5.3% ± 11.2% resulting in a valine to alanine substitution at amino acid position p.2214.

Following base editing, we next confirmed the restoration of PC1 expression by immunoblotting, which showed recovery of PC1 protein levels (**Fig. 2d**). In addition to rescuing PC1 expression, base editing significantly reduced the expression of the endoplasmic reticulum (ER) stress marker sXBP1, upregulated in unedited controls. Densitometric analysis indicated a marked improvement in PC1 levels (**Fig. 2e**) and a significant reduction in sXBP1 (**Fig. 2f**), demonstrating that base editing not only restored the protein expression but also mitigated the associated cellular stress response induced by the pathogenic variant.

### *In vivo* correction of *Pkd1* c.6646C>T (R2216W)

Next, we aimed to evaluate whether base editing can correct the pathogenic *Pkd1* c.6646C>T (R2216W) variant *in vivo* and potentially halt or reverse the phenotypic manifestations of ADPKD. Besides the primary kidney manifestation, cystic liver disease presents a major therapeutic challenge in many cases of ADPKD. Therefore, we selected an ADPKD model that develops both kidney and liver cysts. Importantly, this model enables the evaluation of base editing for ADPKD independently of kidney delivery constraints, as liver delivery is well-established. *Pkd1*^*R2216W/fl*^;*UBC-Cre-ERT2* mice were treated with tamoxifen for 14 days starting at P28, leading to Cre-mediated excision of the floxed allele and resulting in a marked cystic liver phenotype and a mild cystic kidney phenotype (**Fig. 3a**). At P49, these mice were injected intraperitoneally with AAVs at 8×10^11^ viral genomes (VG) per mouse carrying base editing components, followed by analysis at P126. We selected a dual AAV8 approach delivering a split-intein ABEmax^25^ together with sgRNA_45 (**Fig. 3b**). Macroscopic inspection revealed a significant reduction in liver size in ABEmax-treated mice compared to untreated controls, while kidney size appeared similar between the groups. This finding confirms the liver-targeted phenotype of our model and suggests that base editing has a substantial impact on the overall phenotype (**Fig. 3c**). Histological analysis of untreated mice revealed extensively cystic liver tissue affecting all segments and disrupting tissue architecture, whereas base editor-treated mice exhibited only mild phenotypic changes (**Fig. 3d**,**e**). While liver-to-body weight ratios showed no significant differences between groups (**Fig. 3f**), cystic indices were significantly lower in ABEmax-treated mice compared to untreated controls (**Fig. 3g**), highlighting the potential of base editing to reduce cystic burden in ADPKD. Due to the mild cystic kidney phenotype in control mice and the liver tropism of AAV8, kidney-to-body weight ratios did not differ between the groups (data not shown). Next, we explored which editing efficiencies correlated with the observed phenotypic differences. In liver tissue, we observed a mean on-target editing of 5.0% ± 2.6%, compared to 0.5% ± 0.1% in non-edited control mice (**Fig. 3h**), indicating that an overall low editing efficiency is sufficient to modify the cystic liver phenotype. Importantly, no unwanted bystander mutations were detected at A10 or A13 within the editing window. Finally, we examined sgRNA-dependent off-target editing. *In silico* prediction using Cas-OFFinder identified two sites for the human c.6658C>T (R2220W) variant with one and two mismatches, all identified to be located in *PKD1* pseudogenes, and 31 sites with three mismatches, all without RNA or DNA bulges. For the mouse variant c.6646C>T (R2216W), one site with one mismatch was found, and no two-mismatch sites and 21 three-mismatch sites were identified. Targeted amplicon sequencing of the top six predicted off-target sites showed no editing in the liver, indicating precise base editing (**Supplementary Fig. 1a-c**)^31^.

**Fig. 3:**
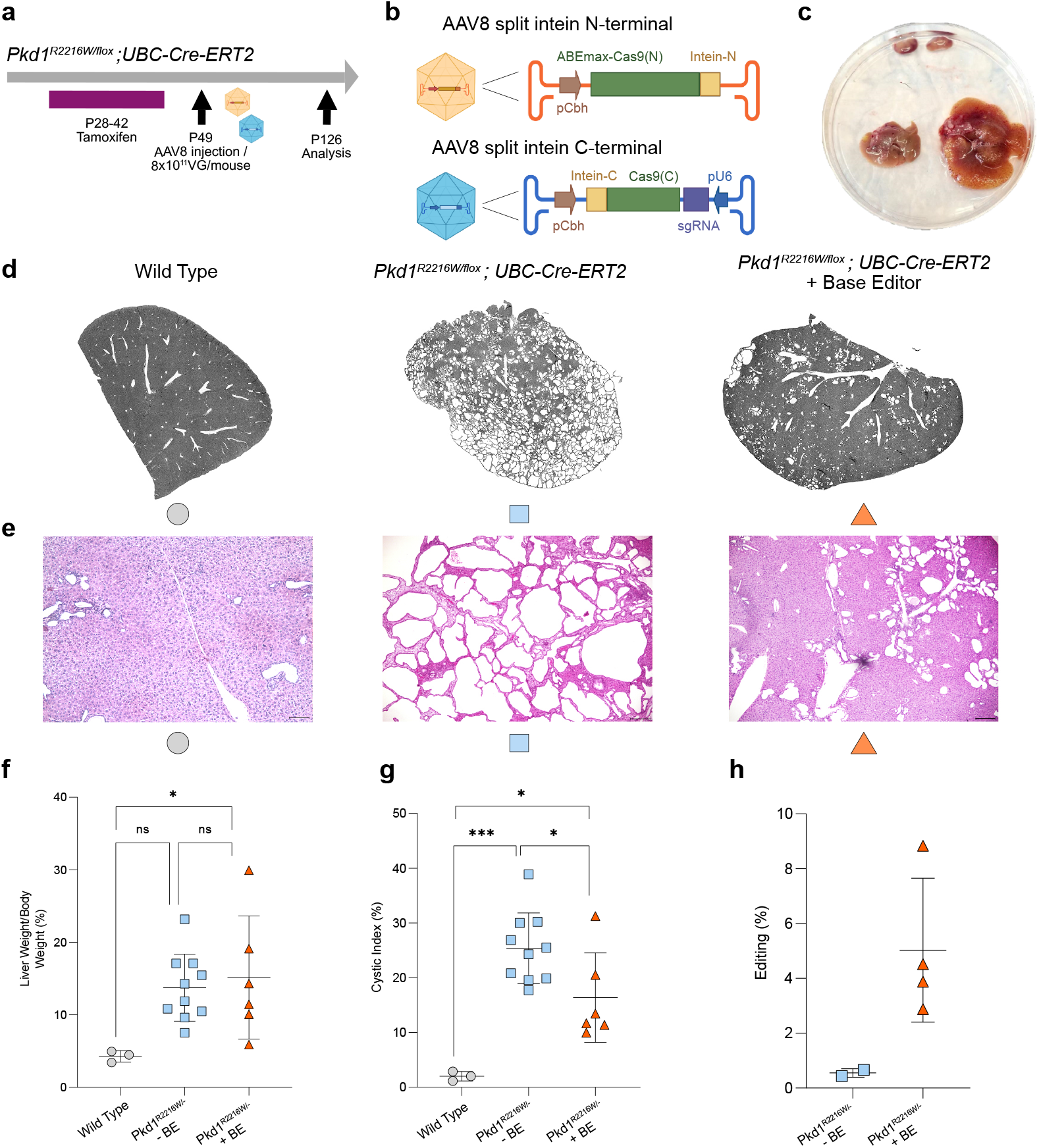
*In vivo* base editing reduces liver cysts in the RW knock-in model. **a,** Schematic illustrating the experimental workflow of tamoxifen mediated Cre expression, AAV8 delivered base editing and analysis at the indicated postnatal (P) days. **b**, Schematic illustrating split-intein based AAV8 delivery of ABEmax. **c**, Macroscopic view of mouse kidney (upper part) and liver tissue (lower part). Left: base editor treated, right: untreated control. **d**, Microscopic overview images of liver tissue from the indicated conditions. **e**, Hematoxylin and eosin (H&E) stain of the indicated conditions. **f**,**g** Quantification of liver to body weight ratios (**f**) and liver cystic indices (**g**) between wild-type mice (n=3), untreated RW knock-in mice (n=10) and base editor (BE) treated RW knock-in mice (n=6). **h**, Quantification of editing efficiency in gDNA isolated from the liver of the untreated RW knock-in mice (n=2) and base editor (BE) treated RW knock-in mice (n=4). **f**,**g**,**h**, data points represent independent biological replicates with means ± SD.

## Discussion

This study presents the first *in vivo* evidence of using base editing to correct pathogenic polycystin-1 variants in ADPKD, providing promising results for the potential reversal of cystic liver disease. Our findings demonstrate that base editing can be applied broadly to human *PKD1* mutations, with correction efficiencies ranging from 30% to 60% for most variants in HEK293T cells.

The restoration of PC1 expression in *Pkd1 R2216W/–* cells after base editing provides further validation of ABE’s potential to restore the normal function of proteins compromised by genetic mutations. The reduction of the endoplasmic reticulum (ER) stress marker sXBP1 further indicates that base editing can alleviate cellular stress associated with pathogenic *PKD1* variants, an important factor in disease progression. Together, these results suggest that ABE not only corrects the underlying genetic defect but also positively impacts disease-related molecular pathways.

Our *in vivo* experiments, conducted in *Pkd1*^*R2216W/fl*^;*UBC-Cre-ERT2* mice, demonstrated a significant reduction in liver cystic burden following a single administration of AAV8-ABE, providing strong initial evidence for the therapeutic potential of base editing in ADPKD. Notably, the editing efficiency required to improve the cystic liver phenotype *in vivo* was comparatively low, underscoring the overall feasibility of this approach. Although the mouse model and the delivery vehicles used here were not designed to investigate and target the kidney phenotype, we observed a trend toward a reduced kidney cyst burden in some base editor treated kidneys (data not shown). Along with recently reported AAV serotypes^32^, this may support the feasibility of targeting renal disease in ADPKD, which represents the major cause of morbidity in affected patients.

Our data suggest that base editing can be used to correct pathogenic mutations in a complex, multiorgan genetic disease. Unlike conventional treatments such as Tolvaptan, which slow disease progression but do not address the underlying genetic cause, ABEs offers the possibility of potentially reversing disease phenotypes by maintaining or restoring normal protein function at the genetic level. The significant reductions in liver cyst burden observed in our study further highlight the potential of base editing to treat extrarenal complications of the disease, which significantly impact morbidity and mortality but remain unaddressed by Tolvaptan, the only licensed medical treatment to date.

While the current study demonstrates the feasibility and therapeutic potential of ABE in ADPKD, several important questions remain. Optimization of delivery methods, particularly to enhance editing efficiency in kidney tubular epithelial cells, will be crucial. Additionally, further work is needed to assess long-term outcomes, genomic stability and potential off-target effects associated with AAV8-ABE delivery. While the overall low *in vivo* editing efficiency was sufficient to alter the hepatic phenotype, improvements in base editors are likely to enhance editing efficiency and phenotypic rescue. Notably, we selected ABEmax due to its lowest degree of bystander editing; however, more active base editors, such as ABE8.20m and ABE8e, are likely to increase editing efficiency, as suggested by our *in vitro* data (**Fig. 2a**).

In conclusion, this study provides initial evidence that adenine base editing is a promising therapeutic strategy for ADPKD. By successfully correcting *PKD1* mutations *in vitro* and *in vivo*, we have demonstrated the potential of base editing to address the genetic root of the disease and reverse cystic phenotypes in the liver. Future work will focus on optimizing delivery to the kidney, increasing editing efficiency, and exploring the full therapeutic potential of ABE in preventing and treating ADPKD.

## Acknowledgements

We thank the Viral Core Facility (VCF) of the Charité – Universitätsmedizin Berlin for producing AAVs. We thank Lonnette Diggs in the George M. O’Brien Kidney Center at Yale (P30 DK079310) for BUN and creatinine measurements.

## Funding Support

MMK was supported by the Emmy Noether Programme (grant no. KA5060/1-1) of the German Research Foundation (DFG) and is a participant in the BIH Charité Clinician Scientist Program funded by the Charité— Universitätsmedizin Berlin and the Berlin Institute of Health at Charité (BIH). AI was supported by the Add-on Fellowship for Interdisciplinary Life Science from the Joachim Herz foundation. JH receives funding from the German Research Foundation (DFG; project number HA 6908/4-1, HA 6908/7-1, HA 6908/8-1, HA 6908/12-1). JH and KUE are members of the European Reference Network for Rare Kidney Diseases (ERKNet). VK is a participant in the BIH Charité Clinician Scientist Program funded by the Charité— Universitätsmedizin Berlin and the Berlin Institute of Health at Charité (BIH) and supported by the Else-Kröner Fresenius Stiftung (Else-Kröner Memorial Grant). RB, FH, and and SF were supported by a DOD Investigator-Initiated Research Award to SF (GR114977).

**Fig. S1.**
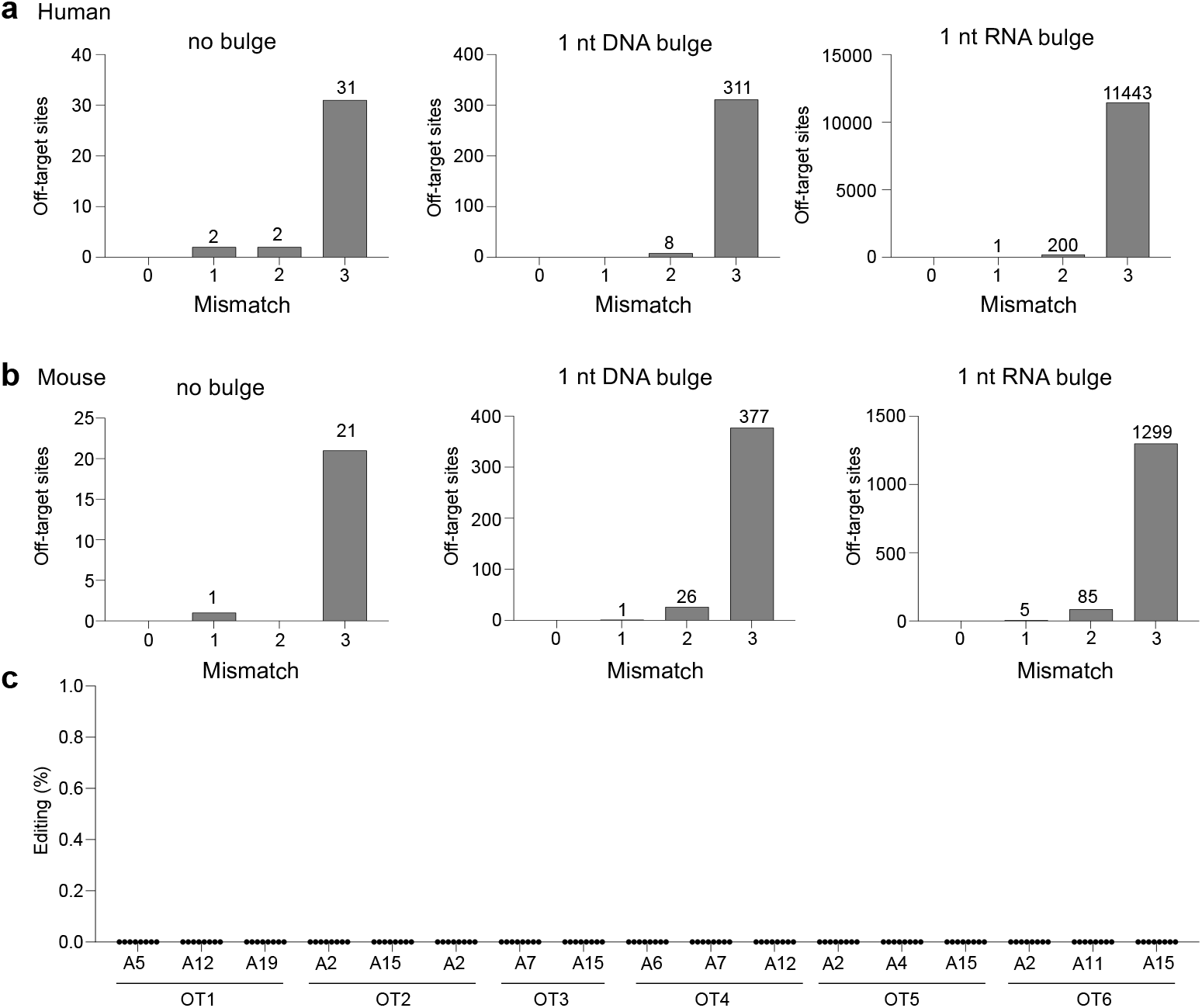
Off-target analysis for RW correcting human and mouse sgRNAs. **a,** Number of CasOFFinder annotated putative off-target sites in the human genome for sgRNA_20 (for SpCas9 and a NGG PAM). Predicted off-target sites for one or two mismatches (without bulges) were identified within *PKD1* pseudogenes. **b**, Number of CasOFFinder annotated putative off-target sites in the mouse genome for sgRNA_45 (for SpCas9 and a NGG PAM). The one site predicted for one mismatch (no bulge) marks the target site. **c**, *In vivo* editing efficiency (y-axis) for the top 6 predicted mouse off-target sites (OT) as determined by targeted amplicon sequencing of gDNA isolated from liver tissue at all possible adenines (A).

## References

1. Bergmann, C. et al. Polycystic kidney disease. Nat Rev Dis Primers 4, 1–24 (2018).

2. Cornec-Le Gall, E., Alam, A. & Perrone, R. D. Autosomal dominant polycystic kidney disease. Lancet 393, 919–935 (2019).

3. Hopp, K. et al. Functional polycystin-1 dosage governs autosomal dominant polycystic kidney disease severity. J Clin Invest 122, 4257–4273 (2012).

4. Lantinga-van Leeuwen, I. S. et al. Lowering of Pkd1 expression is sufficient to cause polycystic kidney disease. Hum Mol Genet 13, 3069–3077 (2004).

5. Qian, F., Watnick, T. J., Onuchic, L. F. & Germino, G. G. The molecular basis of focal cyst formation in human autosomal dominant polycystic kidney disease type I. Cell 87, 979–987 (1996).

6. Dong, K. et al. Renal plasticity revealed through reversal of polycystic kidney disease in mice. Nat Genet 53, 1649–1663 (2021).

7. Lakhia, R. et al. PKD1 and PKD2 mRNA cis-inhibition drives polycystic kidney disease progression. Nat Commun 13, 4765 (2022).

8. Kurbegovic, A., Pacis, R. C. & Trudel, M. Modeling Pkd1 gene-targeted strategies for correction of polycystic kidney disease. Mol Ther Methods Clin Dev 29, 366–380 (2023).

9. Komor, A. C., Kim, Y. B., Packer, M. S., Zuris, J. A. & Liu, D. R. Programmable editing of a target base in genomic DNA without double-stranded DNA cleavage. Nature 533, 420–424 (2016).

10. Gaudelli, N. M. et al. Programmable base editing of A•T to G•C in genomic DNA without DNA cleavage. Nature 551, 464–471 (2017).

11. Tong, H. et al. Programmable A-to-Y base editing by fusing an adenine base editor with an N-methylpurine DNA glycosylase. Nat Biotechnol 41, 1080–1084 (2023).

12. Kurt, I. C. et al. CRISPR C-to-G base editors for inducing targeted DNA transversions in human cells. Nat Biotechnol 39, 41–46 (2021).

13. Koblan, L. W. et al. In vivo base editing rescues Hutchinson-Gilford progeria syndrome in mice. Nature 589, (2021).

14. Musunuru, K. et al. In vivo CRISPR base editing of PCSK9 durably lowers cholesterol in primates. Nature 593, 429–434 (2021).

15. Ryu, S. M. et al. Adenine base editing in mouse embryos and an adult mouse model of Duchenne muscular dystrophy. Nature Biotechnology 36, 536–539 (2018).

16. Villiger, L. et al. Treatment of a metabolic liver disease by in vivo genome base editing in adult mice. Nat Med 24, 1519–1525 (2018).

17. Koblan, L. W. et al. Improving cytidine and adenine base editors by expression optimization and ancestral reconstruction. Nature Biotechnology 36, 843–848 (2018).

18. Gaudelli, N. M. et al. Directed evolution of adenine base editors with increased activity and therapeutic application. Nature Biotechnology 38, 892–900 (2020).

19. Anzalone, A. V. et al. Search-and-replace genome editing without double-strand breaks or donor DNA. Nature (2019) doi:10.1038/s41586-019-1711-4.

20. Alves, C. R. R. et al. Optimization of base editors for the functional correction of SMN2 as a treatment for spinal muscular atrophy. Nat Biomed Eng 8, 118–131 (2024).

21. Richter, M. F. et al. Phage-assisted evolution of an adenine base editor with improved Cas domain compatibility and activity. Nature Biotechnology 38, 883–891 (2020).

22. Walton, R. T., Christie, K. A., Whittaker, M. N. & Kleinstiver, B. P. Unconstrained genome targeting with near-PAMless engineered CRISPR-Cas9 variants. Science 368, 290–296 (2020).

23. Clement, K. et al. CRISPResso2 provides accurate and rapid genome editing sequence analysis. Nature Biotechnology 37, 224–226 (2019).

24. Mátés, L. et al. Molecular evolution of a novel hyperactive Sleeping Beauty transposase enables robust stable gene transfer in vertebrates. Nat Genet 41, 753–761 (2009).

25. Levy, J. M. et al. Cytosine and adenine base editing of the brain, liver, retina, heart and skeletal muscle of mice via adeno-associated viruses. Nature Biomedical Engineering 4, 97–110 (2020).

26. Krappitz, M. et al. XBP1 Activation Reduces Severity of Polycystic Kidney Disease due to a Nontruncating Polycystin-1 Mutation in Mice. J Am Soc Nephrol 34, 110–121 (2023).

27. Ma, M., Tian, X., Igarashi, P., Pazour, G. J. & Somlo, S. Loss of cilia suppresses cyst growth in genetic models of autosomal dominant polycystic kidney disease. Nat Genet 45, 1004–1012 (2013).

28. Shibazaki, S. et al. Cyst formation and activation of the extracellular regulated kinase pathway after kidney specific inactivation of Pkd1. Hum Mol Genet 17, 1505–1516 (2008).

29. Rees, H. A. & Liu, D. R. Base editing: precision chemistry on the genome and transcriptome of living cells. Nat Rev Genet 19, 770–788 (2018).

30. Vujic, M. et al. Incompletely penetrant PKD1 alleles mimic the renal manifestations of ARPKD. J Am Soc Nephrol 21, 1097–1102 (2010).

31. Bae, S., Park, J. & Kim, J.-S. Cas-OFFinder: a fast and versatile algorithm that searches for potential off-target sites of Cas9 RNA-guided endonucleases. Bioinformatics 30, 1473–1475 (2014).

32. Furusho, T. et al. Enhancing gene transfer to renal tubules and podocytes by context-dependent selection of AAV capsids. Nat Commun 15, 10728 (2024).

